# *Frem2* Knockout Mice Exhibit Fraser Syndrome Phenotypes and Neonatal Lethality Due to Bilateral Renal Agenesis

**DOI:** 10.1101/2024.10.28.620501

**Authors:** Rubina G. Simikyan, Xinyuan Zhang, Olga Strelkova, Nathan Li, MengYu Zhu, Andreas Eckhard, Petr Y. Baranov, Xudong Wu, Lauren Richey, Artur A. Indzhykulian

**Affiliations:** Department of Otolaryngology Head and Neck Surgery, Mass Eye and Ear, Harvard Medical School, Boston, MA, United States; Tufts Comparative Medicine Services, Tufts University, Boston, MA, United States; Department of Neurobiology, Harvard Medical School, Boston, MA, United States; Department of Ophthalmology, Massachusetts Eye and Ear, Harvard Medical School, Boston, MA, United States

**Keywords:** Fraser syndrome, FREM2, embryogenesis, bilateral renal agenesis, cryptophthalmos, syndactyly

## Abstract

Fraser syndrome is a rare autosomal recessive disorder characterized by multiple congenital malformations, including cryptophthalmos, syndactyly, and renal agenesis, which can lead to severe complications beginning at the embryonic stage. Mutations in genes encoding extracellular matrix proteins such as FRAS1, FREM1, FREM2, and the associated trafficking protein GRIP1, are implicated in Fraser syndrome. These proteins are critical for maintaining epithelial integrity during embryogenesis, with deficiencies leading to tissue detachment and blistering phenotypes in mouse models. The FREM2 protein is a single-pass membrane protein of 3169 amino acids. While *Frem2-*deficient mouse models encoding missense variants found in patients, or a truncated FREM2 protein product were previously reported, it has not been studied in a constitutive knockout (KO) mouse model.

Here, we developed constitutive *Frem2-KO* mice exhibiting neonatal lethality, mainly due to bilateral renal agenesis, along with blood-filled blisters, cryptophthalmos, and syndactyly. Only one mouse survived to adulthood exhibiting unilateral renal agenesis and Fraser syndrome-like phenotypes. These findings confirm FREM2’s crucial role in the development of the kidneys, skin, and eyes and provide an animal model for further studies of FREM2-related developmental disorders.

## INTRODUCTION

Fraser syndrome is a rare autosomal recessive disorder characterized by developmental malformations evident before birth [1]. Individuals affected may exhibit various features such as cryptophthalmos (fused eyelids), syndactyly (fused fingers and toes), and unilateral or bilateral renal agenesis (failure of kidney development), alongside respiratory and ear abnormalities [2]. Despite its rarity, Fraser syndrome may contribute to early-term miscarriages, mainly due to renal or pulmonary complications at the embryonic stage [1]. Patients without life-threatening phenotypes can survive into adulthood and may utilize surgical interventions to resolve skin malformations such as syndactyly.

FRAS1, FREM1, and FREM2 are structurally similar proteins shown to serve as important components of the extracellular matrix (ECM) protein complex [3]. These proteins are predominantly localized in the skin within the *sublamina densa*, a component of the basement membrane zone between the epidermis and dermis of the skin, playing a critical role in preserving epithelial-mesenchymal integrity. GRIP1 is involved in trafficking the ECM proteins to their correct location, making it a crucial protein for mediating organ morphogenesis. Although it is not included in the ECM protein complex, it was suggested to play an important role in maintaining the structural integrity of tissues [4]. Pathogenic variants in the *FRAS1, FREM1, FREM2*, and *GRIP1* genes cause epithelial detachment at the level of the *sublamina densa* [3, 4].

Previous studies reported that pathogenic variants in *FRAS1, FREM2, and GRIP1* are causative of Fraser syndrome, however pathogenic variants in *FREM1* alone do not induce the disorder [5]. Nonetheless, pathogenic variants in these genes result in similar phenotypes. The severity of these phenotypes in patients, caused by deleterious pathogenic variants or the absence of ECM proteins and GRIP1 highlights their requirement for assembling basement membranes across critical organs such as the skin, kidneys, testes, and trachea during embryogenesis [3]. Mouse models have proven an excellent tool for investigating the roles of key proteins in embryonic development and explaining their phenotypic correlations. Although FRAS1, FREM1, and FREM2 are predicted to be structurally similar, genetic studies in mouse models indicate that they are functionally non-redundant; loss of any component destabilizes the entire basement membrane complex, suggesting that these proteins are likely to form a complex and reciprocally stabilize one another at the epithelial basement membrane [6].

Previously reported mouse models carrying mutations in this group of genes have displayed similar blistering phenotypes, often referred to collectively as “bleb” mutants [3]. Loss of function in *Grip1* results in blistering over the eyes, mimicking the phenotype of the *eye blebs* (*eb*) mutant strain [5]. Mutations in *Fras1* in mice produce paw *blebs* (*bl*) in the *Fras1*^*bl/bl*^ mice, while *Frem1* mutations result in *head blebs* (*heb*) in mice, often presenting as head blisters and malformed or absent eyes at birth [7]. The Myelencephalic bleb (*my*) mouse line, identified as early as the 1920s and currently available from The Jackson Laboratory, has been linked to the *Frem2* gene, although the underlying genetic lesion remains undefined as no coding region mutations were found in *Frem2* [6]. Reduced FREM2 expression was observed in *Frem2*^*my/my*^ embryos, but it remains unclear whether this results from cis-regulatory disruption or transcript instability, and the allele has not been confirmed as a functional null [6]. Another strain, carrying the *Frem2-my*^*UCL*^ allele caused by a random transgene insertion event, was reported to localize to the *Frem2* gene; however, the precise mutation has not been identified, the transgene was not detected, and no coding region mutations were found in *Frem2* [8]. *Fras1*- and *Grip1*-knockout mouse models also show early-onset blistering, detectable by embryonic day 12-13, and both exhibit embryonic lethality [9, 10].

Mutations in *Frem2* have been associated with a spectrum of Fraser syndrome-like phenotypes, including cryptophthalmos, epithelial blebbing, blood-filled blisters, renal agenesis, and bony syndactyly [8, 11]. Among the previously reported *Frem2* mouse models are the compound heterozygous *Frem2*^*R725X/R2156W*^ mice, which carry patient-derived variants associated with cryptophthalmos, and the *Frem2*^*my-F11*^ mice which harbor a nonsense mutation resulting in a premature stop codon in Exon 5 of 24 [11, 12]. Most, if not all, of these models carry variants predicted to produce truncated FREM2 protein or single amino acid substitutions, both of which may retain partial function (**Supplementary Table 1**). These alleles are often associated with milder phenotypes, with the deficit limited to specific cell types or tissues, such as the cryptophthalmos phenotype of the *Frem2*^*R725X/R2156W*^ mouse model [11]. Therefore, a constitutive *Frem2* knockout (*KO*) mouse model is needed to fully elucidate FREM2’s role in development and to generate a more complete phenotypic picture of Fraser syndrome.

To assess whether the absence of FREM2 recapitulates the reported phenotypes in *Frem2* mutants, or perhaps, results in a more exacerbated phenotype, we developed and characterized a constitutive *Frem2-KO* mouse model. Upon anatomical and histological analysis, we found that the *Frem2-KO* mice exhibit neonatal mortality which we associate with bilateral renal agenesis. In addition, the *Frem2-KO* fetuses develop blood-filled blisters on the eyes and paws that progress into hemorrhages and missing eyelids. A single *Frem2-KO* mouse survived to adulthood and displayed unilateral renal agenesis while exhibiting syndactyly, cryptophthalmos, and microphthalmia. Our findings confirm FREM2’s role as an important protein for the formation of the skin, kidneys, and eyes. This *Frem2-KO* mouse model provides a valuable tool for further, more detailed prenatal investigation of its critical roles in organ development and underlying phenotypes resulting from its absence.

## RESULTS

### Genetic analysis of Frem2-KO mouse model

Mouse ES cell clones were purchased from the European Conditional Mouse Mutagenesis Program and used to generate the *Frem2*^*tm1a(EUCOMM)Hmgu*^ mouse line carrying the knockout-first allele with conditional potential (F2KCR allele) (**Fig. 1A**). First, the FRT-flanked Neo cassette was excised by crossing with a FLP deleted strain to generate the conditional *Frem2*^*fl/fl*^ allele. Although the generated mouse line was designed to carry a floxed *Frem2* allele to enable cell-type specific FREM2 functional studies (**Fig. 1A**), following several breeding steps with pan-Cre and tissue-specific Cre lines (see *Methods*), we were unable to generate adult homozygous *Frem2*-floxed mice. Upon further investigation of the mouse genetics, and the sequencing results of the insertion part of the floxed-*Frem2* mouse, we discovered mutations within the targeted insertion site. These included two in-frame insertions (27 bp and 30 bp) and one in-frame deletion (9 bp) (**Fig. 1A**). Subsequently, we analyzed the open reading frame in exon 1 of the floxed *Frem2* allele. We identified a premature stop codon (**Fig. 1B**) within exon 1 in the Floxed-*Frem2*, which likely results in a truncated or missing FREM2 protein, converting the floxed-*Frem2* allele by design into a constitutive null allele. All animals evaluated in this study were obtained from *Frem2*^*+/-*^ x *Frem2*^*+/-*^ crosses.

**Figure 1.**
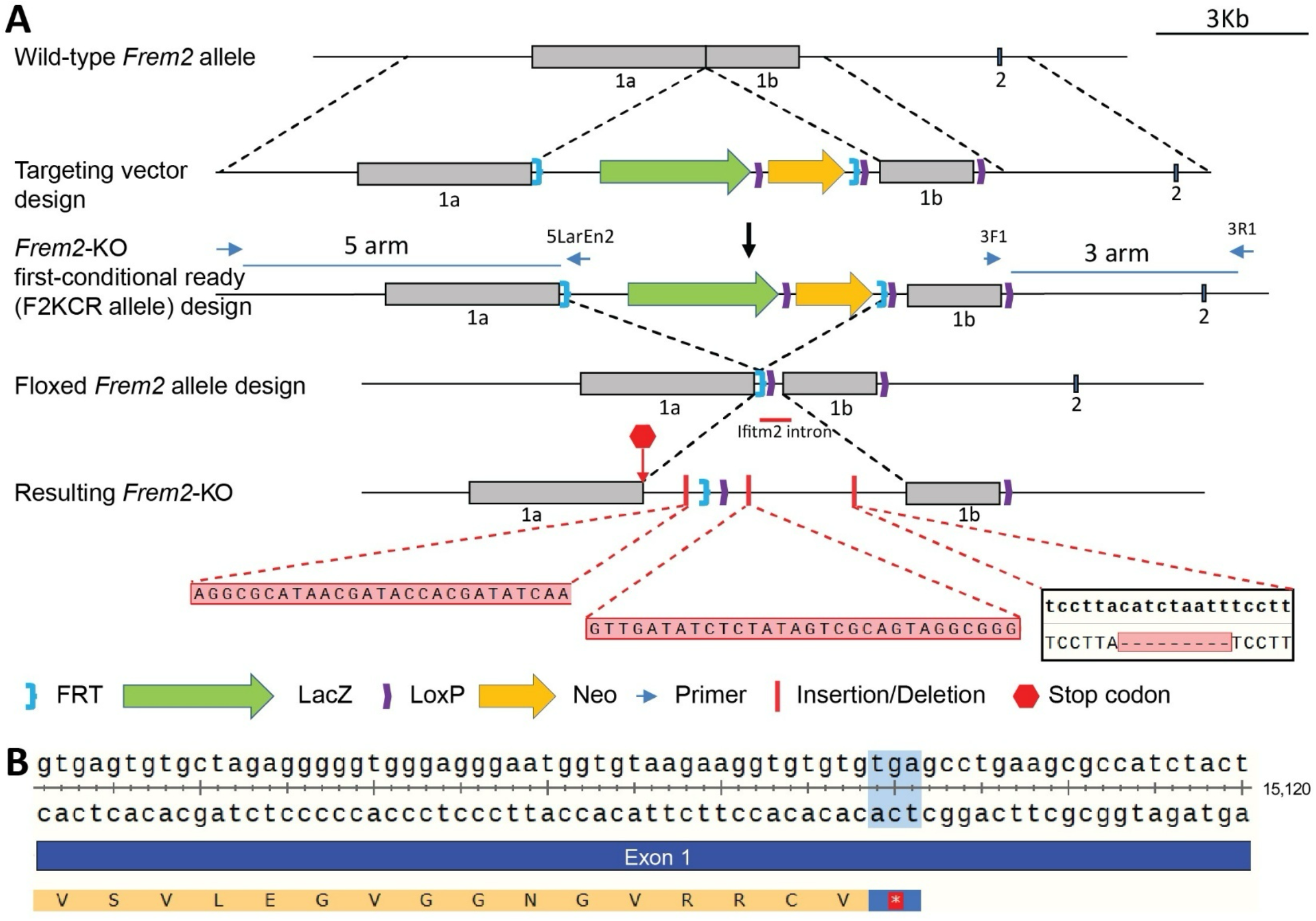
*Frem2-KO* mouse design strategy. (***A***) ES cells were with a *Frem2-KO*-first conditional ready allele was used to generate the Floxed-*Frem2* mouse line. The floxed allele shows the FLP and FRT system inserted into exon 1, dividing it into 1a and 1b regions. The conditional-KO mouse displays the in-frame mutations (shown as a red line) and their sequences: two insertions (27 bp and 30 bp) and one deletion (9 bp). (***B***) A portion of the sequence (15100-15120 bp) for exon 1 of Floxed-*Frem2* is displayed. The premature TGA stop codon sequence is highlighted in blue near the middle of the reading frame.

### Frem2-KO mice have hemorrhagic blisters and skeletal malformations on their paws

Upon our examination of fetuses (E13-16), we immediately observed the skin deficit phenotype on their paws, in agreement with previous studies reporting FREM2’s critical role in epidermis development. Hemorrhagic blisters were observed on the digits of *Frem2-KO* fetuses at E15 and E16 (**Fig. 2A** and **B**) which allowed to phenotype them when compared to *Frem2* wild-type and heterozygote fetuses that appeared normal. To observe the postnatal development of the limbs, newborn *Frem2* pups were collected. When we assessed blood-filled blisters on the digits of newborn *Frem2-KO* pups, they were still present but often appeared dry (**Fig. 2C**, arrowheads).

**Figure 2.**
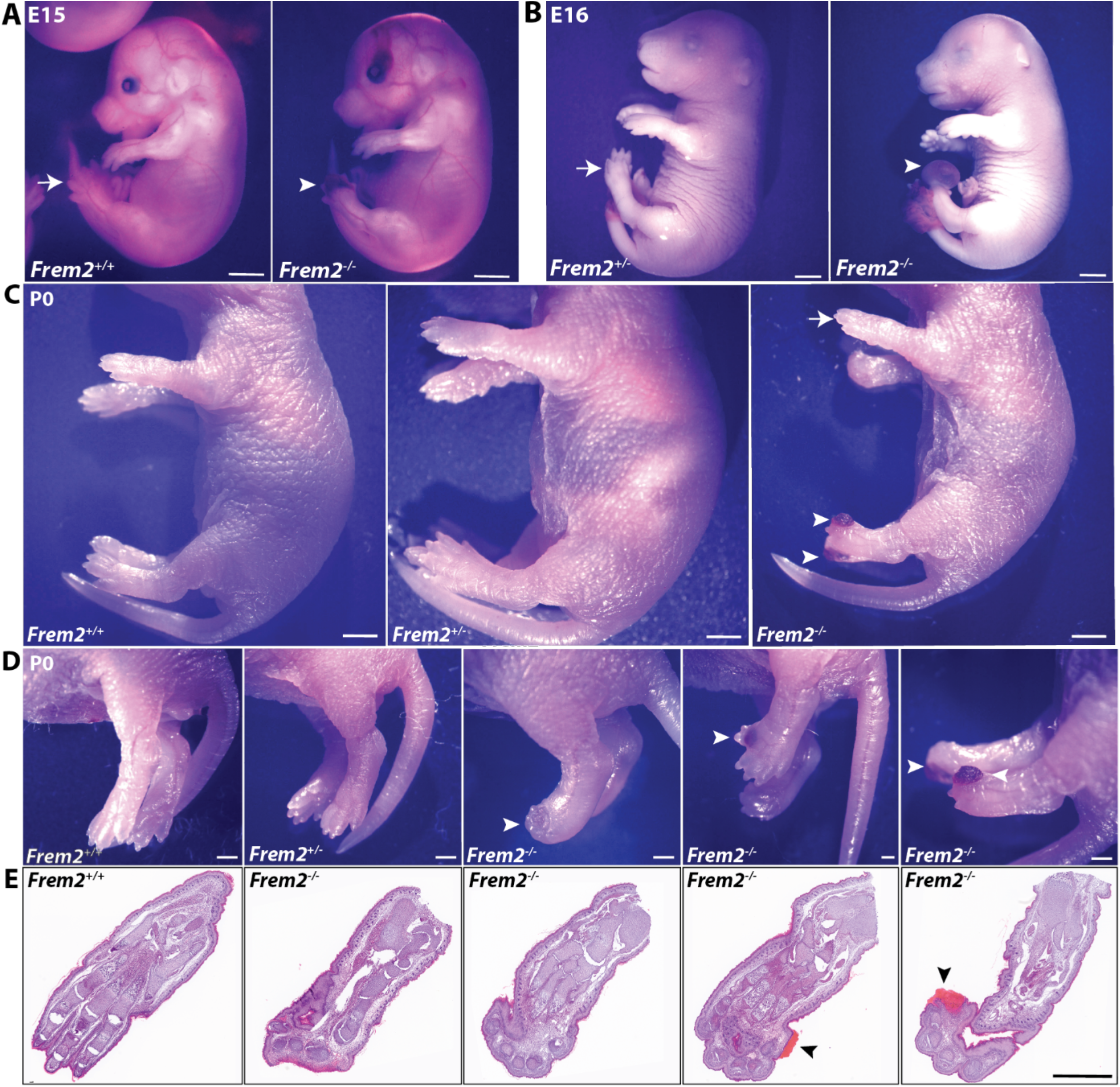
*Frem2-KO* fetuses and newborn pups exhibit hemorrhagic blisters and syndactyly. (***A***) *Frem2*^*+/+*^(wild-type) E15 fetus (*left*) with normal paws (*arrow*) and a *Frem2*^*-/-*^ (KO) littermate (*right*) with a blood-filled blister over the digits (*arrowhead*). (***B***) E16 heterozygous *Frem2*^*+/-*^ fetus (*left*) with normal paws (*arrow*) and E16 *Frem2*^*-/-*^ fetus littermate (*right*) with a large hemorrhagic bleb on the digits (*arrowhead*). (***C***) P0 *Frem2*^*+/+*^ (*left*) and *Frem2*^*+/-*^ (*middle*) have normal paws, and a *Frem2*^*-/-*^ (*right*) has dried blood-filled blisters on both hind paws. (***D***) Newborn *Frem2*^*-/-*^ pups with a malformation of hind limbs, soft-tissue syndactyly, and blood-filled blisters on digits. Reference *Frem2*^*+/+*^ and *Frem2*^*+/-*^ paws showing normal paw morphology. (***E***) Hematoxylin and eosin-stained histology section of paws from wild-type (*Frem2*^*+/+*^) and several knockout (*Frem2*^*-/-*^) mice with limb malformation and blood-filled blisters (*arrowhead*). Scale bars: (*A*)-(*C*): 2 mm, (*D, E*): 1 mm.

In newborn *Frem2-KO* pups, most hind paws revealed soft tissue syndactyly, independent of whether any blisters were observed. Affected paws often also displayed dorsal flexure, an anatomical malformation where the paws curl upwards (**Fig. 2D**). We next assessed the skeletal formation of the paws and other morphological features using histological sections, which were carried out using the same orientation across all samples (**Fig. 2E**). We determined that newborn *Frem2-KO* mice with paws that had blood-filled blisters, syndactyly, or dorsal flexure also exhibited skeletal malformations at the digits, with the skeletal structure around them often compromised. Interestingly, these regions of skeletal malformations typically occurred bilaterally on the hind limbs.

### FREM2 deficiency causes ocular abnormalities in Frem2-KO mice

*Frem2-KO* fetuses consistently exhibited pronounced bubble-like blood-filled blisters over their eyes as early as E13 (**Fig. 3A**) . Coronal histological sections of E15 *Frem2-KO* fetus heads localized the blisters near or within the eyelids (**Fig. 3B** and **C**). In some cases, the eyelids in *Frem2-KO* fetuses were thinned and hemorrhaged (**Fig. 3D** and **E**). Such hemorrhages and blood-filled blisters appeared bilaterally or unilaterally on the eyelids of *Frem2-KO* fetuses (**Fig. 3E, Supplementary Table 2**).

**Figure 3.**
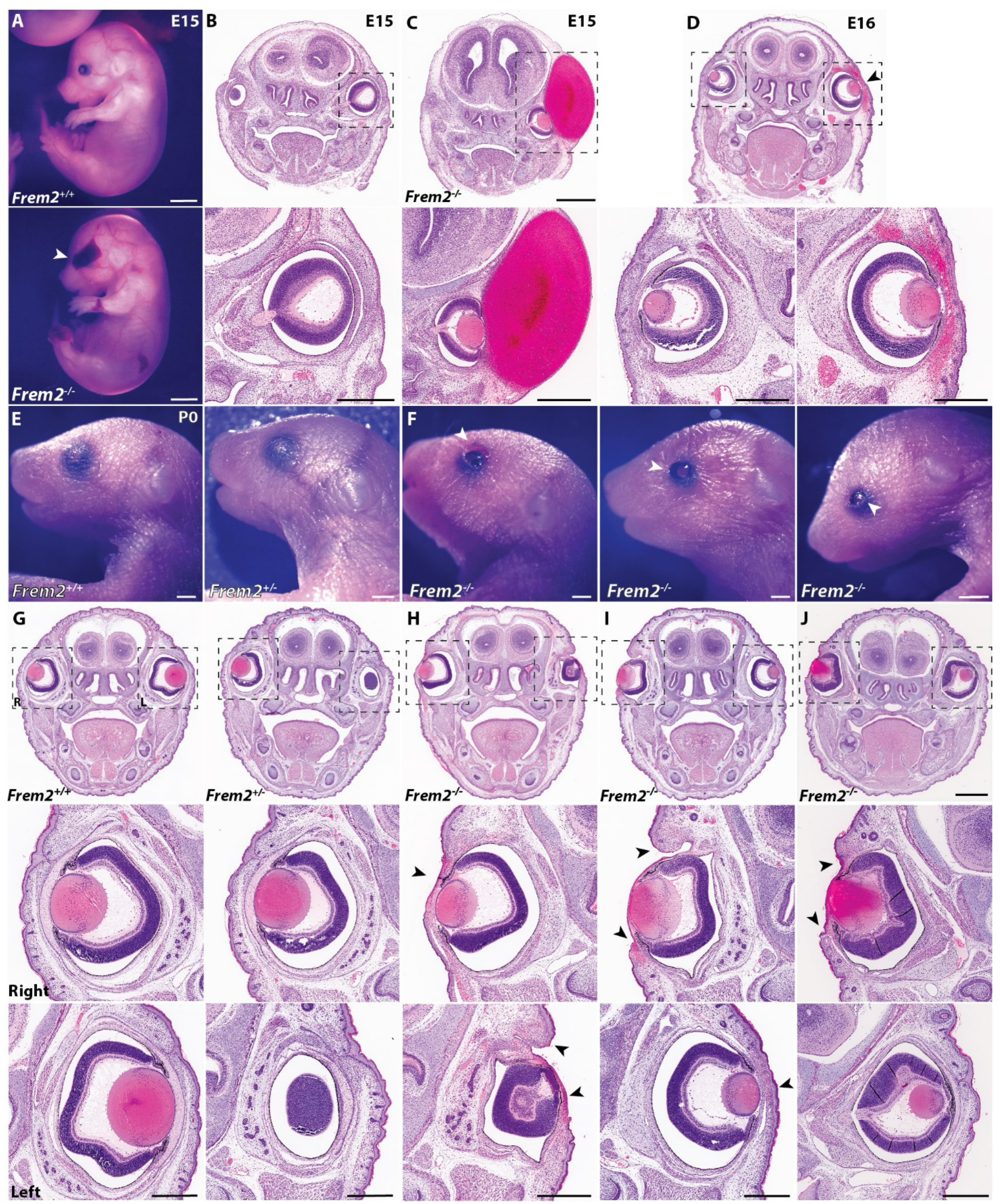
*Frem2-KO* mice lack eyelids and develop blood-filled blisters covering their eyes. (***A***) Gross image of an E15 *Frem2*^*+/+*^ fetus with a normal eye, and a *Frem2*^*-/-*^ fetus (*below*) with a blood-filled blister (*arrowhead*) covering the eye. Hematoxylin and eosin-stained coronal head section of a *Frem2*^*+/-*^ E15 fetus and a higher magnification image below displays normal eyelid morphology. (***C***) Histological section of a *Frem2*^*-/-*^ E15 fetus with a blood-filled blister and a higher magnification image displays the absence of the cornea and eyelid. (***D***) Low and high magnification histology images of a *Frem2*^*-/-*^ E16 fetus with a normal right eyelid and a hemorrhage covering the left eyelid. (***E***) Gross images of *Frem2*^*+/+*^ and *Frem2*^*+/-*^ P0 pups show normal eyelids. (***F***) Gross images of *Frem2*^*-/-*^ P0 pups display missing eyelids (*arrowhead*). (***G***) Histology sections of P0 *Frem2*^*+/+*^ and *Frem2*^*+/-*^ heads with normal eyelids and their corresponding higher magnification images of the right (*R*) and left (*L*) eyes presented below it. (***H***) Histology section of a P0 *Frem2*^*-/-*^ pup with the thinning of the eyelid (*R*) and a missing eyelid with a hemorrhage over the eye (*L*). (***I***) A newborn *Frem2*^*-/-*^ pup with a missing eyelid (*R*) and thinning of the eyelid (*L*). (***J***)*Frem2*^*-/-*^ pup with a missing eyelid (*R*) and a normal contralateral eyelid (*L*). *Scale bars:* (*A*): 2 mm, (*B-G*): 1 mm, (*H-J*): 1 mm.

In newborn *Frem2-KO* pups, however, no blood-filled blisters were observed. Instead, *Frem2-KO* pups predominantly displayed missing eyelids, with some hemorrhages often observed on the skin around the eyes (**Fig. 3F**). The heads of newborn pups were sectioned coronally to assess eye morphology using hematoxylin and eosin staining. In wild-type and heterozygous *Frem2* pups, the epidermal layers of the eyelids were properly formed (**Fig. 3G**). In *Frem2-KO* pups, however, the eyelids were often missing (**Fig. 3H** and **I**) or thinned (**Fig. 3I** and **J**). Periocular hemorrhages and the absence of eyelids were bilateral or unilateral in *Frem2-KO* pups. Blood-filled blisters on the eyelids that occur during embryonic stages, as well as the missing eyelids in newborn *Frem2*-deficient animals, suggest that proper FREM2 expression is critical for the epidermal development of the eyelids.

### Frem2-KO mice die hours after birth

During routine genotyping of *Frem2* litters across several years of the study, no *Frem2-KO* mice were identified. Interestingly, the dam was observed giving birth to pups exhibiting phenotypes of *Frem2-KO* embryos. However, a few hours after birth, the pups were discovered dead in the cage. We thus concluded that *Frem2-KO* pups do not survive after birth, and next sought to investigate the causes of this mortality. Multiple histological sections and post-mortem necropsies were performed in newborn *Frem2-KO* pups from 8 litters. In almost all newborn *Frem2-KO* pups we observed bilateral renal agenesis (**Fig. 4** and **Supplementary Table 2**). Likely a result of the lack of urine production from renal agenesis, *Frem2-KO* pups had empty urinary bladders. We also commonly observed the absence of one or both adrenal glands (**Fig. 4A** and **B**). Parasagittal sections further confirmed the renal agenesis and the empty urinary bladder in newborn *Frem2-KO* pups (**Fig. 4C-F**).

**Figure 4.**
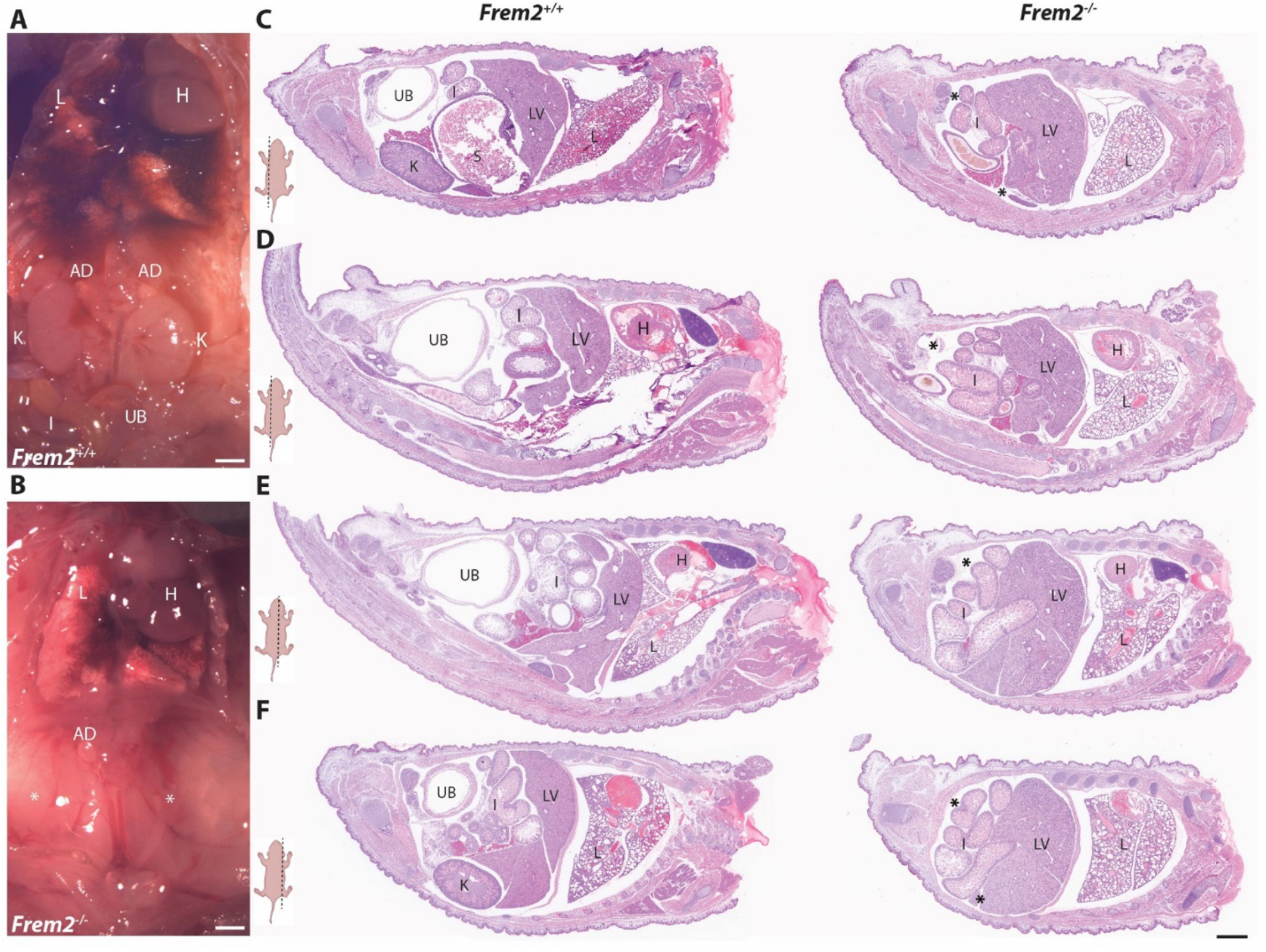
Renal agenesis in *Frem2-KO* pups. Macroscopic images of P0 pups show kidneys (K), adrenal glands (AD), and urinary bladder (UB) present in a *Frem2*^*+/+*^ pup (***A***) and kidneys and inconspicuous urinary bladder in a *Frem2*^*-/-*^ pup (***B***). Hematoxylin and eosin-stained parasagittal sections of a *Frem2*^*+/+*^ (*left column*) and a *Frem2*^*-/-*^ (*right column*) P0 pup show organs present to the left (***C, D***) and right (***E, F***) of the spinal cord. Locations of the deflated urinary bladders are labeled with asterisks. Sections are from similar regions to compare the presence and morphology of organs. Scale bars: (*A, B*); 1 mm, (*C-F*); 3 mm. Labels: AD, adrenal glands; K, kidney; H, heart; LV, liver; LUN, lung; I, intestines; S, stomach; UB, urinary bladder.

We carried out serial transverse histological sections to gain better confidence in our macroscopic results (**Fig. 5** and **Supplementary Movies**). All *Frem2-KO* pups, but one, had bilateral renal agenesis, confirmed by the necropsies or with histological evaluation. A single newborn *Frem2-KO* pup displayed one small but developed kidney, confirming unilateral renal agenesis (**Fig. 5D**). Although the functional capacity of that kidney and chances of that animal’s survival remain unknown, especially considering that a single surviving *Frem2-KO* pup had been observed in previous years. All examined *Frem2-KO* pups also had empty urinary bladders. No major defects in other vital organs, such as the lungs and heart, were seen in *Frem2-KO* pups upon examination with necropsy or histological analysis. Lungs were submerged into phosphate-buffered saline to assess whether they were filled with air and were confirmed to float, suggesting that the animals were breathing before death. We conclude that the likely cause of postnatal mortality of *Frem2-KO* pups is renal agenesis.

**Figure 5.**
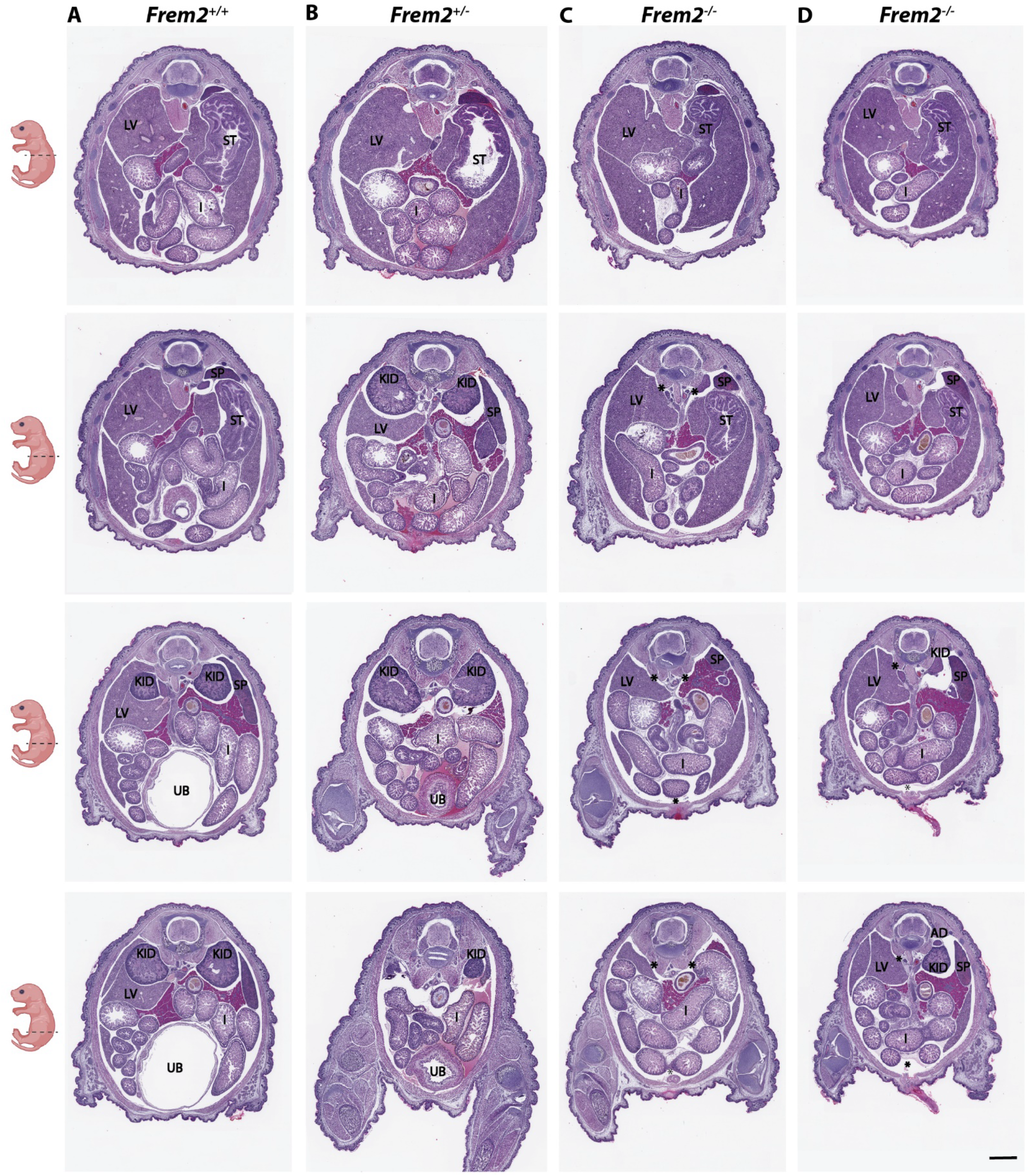
Serial sectioning of newborn (P0) *Frem2* pups reveals renal agenesis in knockout mice. Sections down the columns progress below the diaphragm to the pelvic region from the same pup. Sections across each row correspond to the same region of each pup’s body. Hematoxylin and eosin-stained histology sections from *Frem2*^*+/+*^ (***A***) and *Frem2*^*+/-*^ (***B***) pups exhibit the presence of both kidneys (KID) and a comparable arrangement of organs. *Frem2*^*-/-*^ pups display bilateral and unilateral renal agenesis (***D***). Locations of missing kidneys are labeled with asterisks. Scale bar: (*A-D*); 1 mm. Labels: AD, adrenal glands; KID, kidney; LV, liver; LUN, lung; I, intestines; SP, spleen; UB, urinary bladder.

### A single case of Frem2-KO mouse surviving into adulthood

Over the years of breeding, only one female *Frem2-KO* mouse was identified and survived until adulthood during this study. Although no apparent health conditions were determined and the animal produced two litters, the female was euthanized at the age of 51 weeks due to ulcerative dermatitis, common in older mice. Interestingly, the mouse had displayed physical phenotypes, such as unilateral cryptophthalmos (**Fig. 6A**). The left eye of the mouse was closed, with no visible eyelid crease. The contralateral eye, however, appeared normal. A post-mortem incision through the skin of the closed eyelid, a significantly smaller eyeball was identified and sent for histological analysis along with the contralateral eye. The histological analysis revealed a normally developed right eye and a malformed left eye **(Fig. 6B**). The left eye was dramatically smaller in size, and the retina appeared to have folds, often observed in microphthalmia **(Fig. 6B’**).

**Figure 6.**
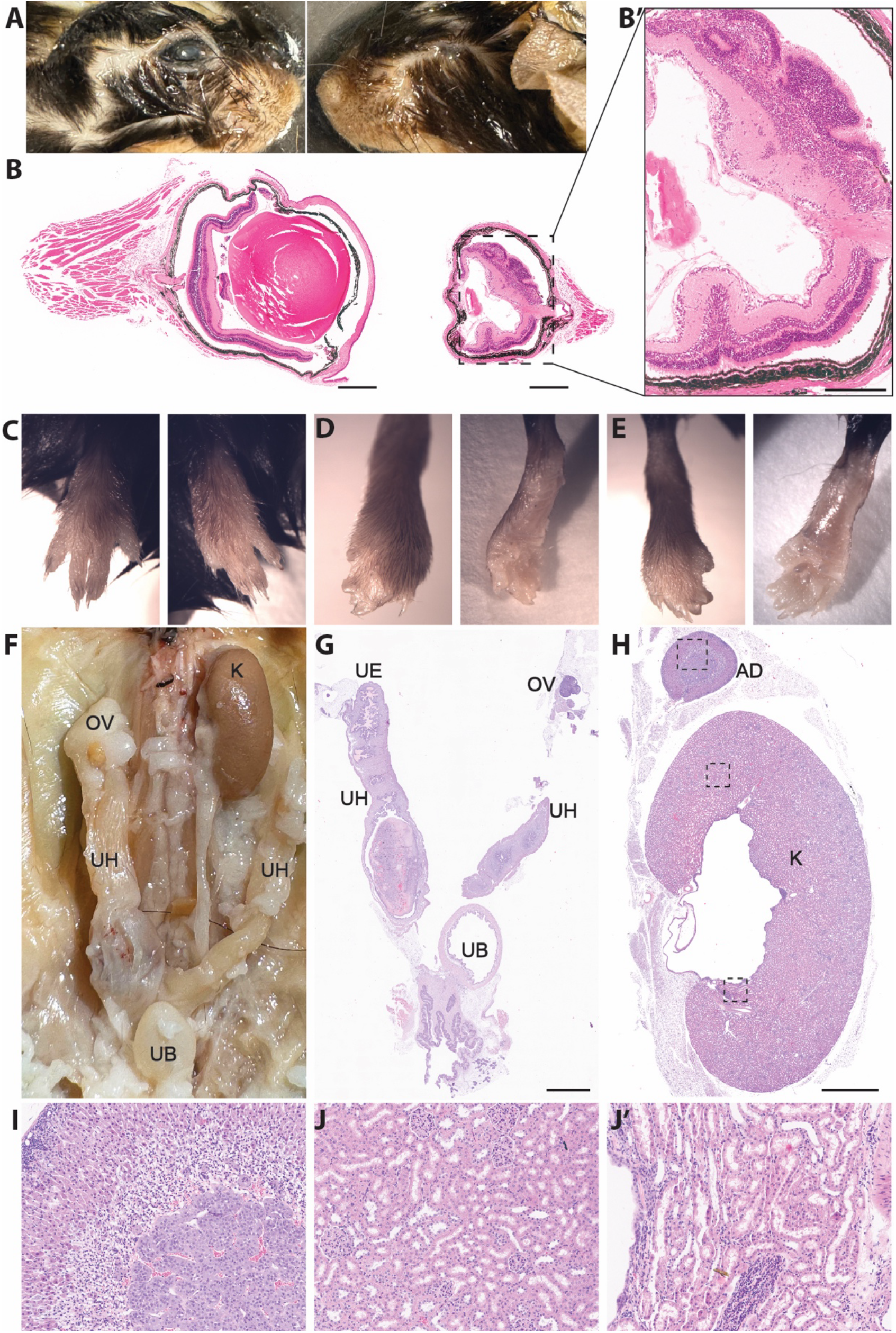
Characterization of the *Frem2-KO* adult mouse. (***A***) Images show the *Frem2*^*-/-*^ adult mouse with a normal right eye and a left eye affected by cryptophthalmos. (***B***) Hematoxylin and eosin-stained histology section through the middle of the normal right eye and the defective left eye. The defective left eye reveals the lens with reduced thickness size and retinal dysplasia. (**B’**) Higher magnification image of defective left eye showing microphthalmia. (***C***) Images of normal front paws, affected right (***D***) and left (***E***) hind paws by syndactyly. (***F, G***) Gross anatomy and histology images of the adult *Frem2*^*-/-*^ mouse show one left kidney (K), ovaries (OV), uterine horns (UH), uterine epithelium (UE), and urinary bladder (UB) present. (***H***) Histology section of the normal left kidney and adrenal gland. (***I-J’***) High-magnification images of the adrenal gland, and different regions of the kidney displaying the tubules, collecting ducts, and nephrons. *Scale bars:* (*B*) 2 mm, (*B’*) 250 um, (*G*) 4 mm, (*H*) 3 mm, (*I-J’*) 500 um.

Consistent with our observations in newborn *Frem2-KO* mice, the front paws of the adult mouse appeared normal (Fig. 6C), while both hind paws were affected by syndactyly (**Fig. 6D** and **E**). The hind paws were also slightly curled, similar to the paws observed in newborns. We next examined the renal system of this adult mouse and found that it had one functional kidney. There was no gross evidence of renal tissue on the right side (**Fig. 6F**). The left kidney was fully attached to the ureter along with the rest of the urinary system. Hematoxylin and eosin histological sections of the left kidney confirmed normal morphology of the adrenal gland and kidney (**Fig. 6H**). Higher magnification images of the histological sections reveal the normal histology of the adrenal gland, as well as of the kidney, containing the unaffected appearance of tubules and nephrons (**Fig. 6I-J’**). The histological evaluation also showed a missing right ureter (**Fig. 6G**). Although missing a kidney, the reproductive functions were not affected in the *Frem2-KO* female, allowing it to breed and produce litters. The renal pelvis was also dilated, according to gross examination. As a result of the functional kidney, the urinary bladder was full (**Fig. 6F** and **G**). Although this female was able to survive with one functional kidney, the *Frem2-KO* adult mouse displayed other prominent Fraser syndrome phenotypes, such as cryptophthalmos and syndactyly (**Fig. 6A, C** and **D**).

## DISCUSSION

In this study, we developed a constitutive knockout (*KO*) *Frem2* mouse model to evaluate the associated phenotypes. Our findings reveal that the absence of the FREM2 protein results in significant Fraser syndrome-like phenotypes, including cryptophthalmos, syndactyly, and blood-filled blisters, observable as early as during the embryonic stage. *Frem2-KO* pups displayed distinct phenotypes that were evident within the litter, even before genotyping was performed. Although *Frem2-KO* mice can survive until birth, they die shortly thereafter likely due to bilateral renal agenesis. To date, only one *Frem2-KO* mouse with unilateral renal agenesis from our animal colony has survived into adulthood. These results highlight the critical role of FREM2 in the development of the epidermis, eyes, skeletal structure, and kidneys.

The extracellular matrix (ECM) is well-known for its role in providing structural support. However, it also plays a crucial role in maintaining tissue integrity by regulating cell proliferation, differentiation, and survival. FRAS1, FREM1, and FREM2 serve as important components of the ECM protein complex, and are structurally similar proteins. Previous studies show that absence of any one of these proteins causes severe phenotypes, suggesting that they are unable to substitute for each other in any major capacity. As previously reported in *Frem2* mutant mice, not only FREM2 but FRAS1 and QBRICK/FREM1 were depleted from the basement membrane zone, suggesting that loss of one complex member leads to depletion of the others, indicating interdependence [6].

The ECM includes basement membranes that form sheets underlying epithelial and endothelial cell layers. FREM2 is localized within the epithelial basement membrane. Previous studies using immunogold labeling have demonstrated the clustered localization of FREM2 within the *sublamina densa* of embryonic skin, highlighting its importance in tissue integrity [3]. Mouse models with a deficiency of ECM proteins, such as FREM2, reveal how disruptions in ECM components can lead to blistering phenotypes, highlighting the intricate relationship between genetic mutations and the assembly of essential basement membranes [13]. Although we do not expect a complete loss of *Frem2* transcript and were unable to confirm a complete loss of protein due to the lack of reliable commercially available anti-FREM2 antibodies, the resulting consistent and severe phenotype observed in *Frem2-KO* mice suggests that this allele is likely to represent a functional null. We acknowledge this limitation to the study and note that custom antibody generation may be necessary for definitive protein-level confirmation, as done in prior studies [6, 14]. Please refer to *Materials and Methods* for a list of antibodies used in this study.

Our observations support previously reported evidence from other studies that use *Frem2* mouse models in reporting that mutations in *Frem2* can cause blood-filled blisters and hemorrhages on the skin. For instance, in a study involving a *Frem2* mouse model of the *my*^*Ucl*^ strain, homozygote mutant mice exhibited epithelial blebbing beginning at E11.5 [8]. By E14, these blebs had progressed into hemorrhagic lesions. Another study focused on embryonic development suggests that the onset of angiogenesis, the process of new blood vessel formation involving the growth and differentiation of endothelial cells, may explain the timing of these hemorrhages [12]. Inadequate adhesion of endothelial cells to adjacent structures, particularly in cells deficient in FREM2 protein, could explain deficits in angiogenesis [12]. Although FREM2 is not directly localized within vascular structures and instead was located in the membrane that lines the blood vessels, its role in stabilizing blood vessels during development appears critical [12]. The absence of FREM2 in surrounding tissues likely contributes to the hemorrhages observed on the skin which aligns with the parallel occurrence of syndactyly and blood-filled blisters observed on the paws and eyes of *Frem2-KO* mice.

The skeletal malformations, syndactyly, and dorsal flexure observed in Frem2-KO mice are unlikely to result from primary defects in bone development, as *Frem2* has not been directly implicated in osteogenesis. This further supports the conclusion that FREM2 is essential not only for vascular stability but also for tissue remodeling and skeletal development during gestation. Supporting this interpretation, studies involving mouse models lacking ubiquitous basement membrane proteins, such as *Nidogen 1* and *Nidogen 2*, have shown that disruption of the ectodermal basement membrane alone is sufficient to cause limb malformations [15], reinforcing the idea that basement membrane integrity is essential for normal limb development. However, the potential relationship between these anomalies remains speculative and warrants further investigation through targeted histological and morphological analyses of skin, bone, or cartilage to elucidate the underlying developmental mechanisms.

Patients with Fraser syndrome who exhibit cryptophthalmos are born with skin covering their eyes or with fused eyelids. Instead in *Frem2*-deficient mice, this phenotype more often presents as the absence rather than the fusing of eyelids. We observed that *Frem2-KO* fetuses developed blood-filled blisters or hemorrhages that cover their eyes and give way to one or both eyelids’ absence at birth. A study on *Frem2* mice carrying the *my*^*F11*^ mutation, which is predicted to result in a truncated protein, indicated that embryonic hemorrhages might lead to localized tissue necrosis, which could explain the absence of eyelids [12]. Given the proposed role of FREM2 in vascular stability, it is likely that the loss of FREM2 during key morphogenic events, like eyelid development, could lead to such hemorrhages. The subsequent blood-filled blisters and hemorrhages likely inhibit normal skin development, causing the observed phenotypes.

Moreover, a study utilizing a compound heterozygous mutation derived from a Fraser syndrome patient to generate mice that mimic the human cryptophthalmos phenotype found significant abnormalities in eyelid development during the critical stages at E13-14 [11]. Specifically, the lower eyelid fold was poorly defined in *Frem2* mutant fetuses, in contrast to wild-type mice, where the grooves of the ectoderm that eventually form the upper and lower eyelids are visible. These mutants exhibited dysplasia and microphthalmia, with reductions in the eye’s axial length and lens size. Eyes affected by microphthalmia typically have a thicker cornea, an absent or severely underdeveloped lens, and a retina prone to folding or filling the vitreous body. These features likely exert pressure on the lens, potentially exacerbated by the presence of blisters or hemorrhages [16]. In our *Frem2-KO* model, we observed microphthalmia where one eye was smaller than the other, as well as retinal folding. In Fraser syndrome patients, these ocular abnormalities often lead to impaired vision. This shows that FREM2 is directly involved in eyelid development and indirectly in the development of the lens, retina and other parts of the eye. While the scRNAseq data suggests that FREM2 is also expressed at early stages of human retinal development [17], in the current study we could not detect early changes in the developing retina.

FREM2 protein was reported to be expressed in the epithelia in the renal cortex, or the outer layer, of mouse kidneys [18]. FREM2 expression begins in the ureteric epithelia of the metanephros at E11.5, with peak expression at the tips of the ureteric buds [8]. In adult mice, FREM2 is strongly expressed in the collecting ducts, proximal convoluted tubules, and arterioles within the kidneys [8]. This expression pattern, along with the development of renal cysts in *Frem2* mutant animals, suggests that FREM2 is essential for maintaining renal integrity [18]. The absence of kidneys in *Frem2-KO* mice that fail to survive postnatally provides additional evidence of FREM2’s critical involvement in renal development. Interestingly, over multiple years of breeding, the only *Frem2-KO* mouse that survived into adulthood had one functional kidney, suggesting that perhaps while FREM2 is critical for renal development, it is not absolutely indispensable. This partial necessity may be due to protein-protein interactions within the ECM protein complex, where other proteins such as FREM1 or FRAS1 may, at least partially, compensate for the loss of FREM2 in the formation of the basement membrane. Although Fraser syndrome is associated with a risk of miscarriage, bilateral renal agenesis is a significant phenotype contributing to this, while patients with unilateral renal agenesis can survive with a single kidney [1]. Since kidneys contribute to the amniotic fluid in humans, by the second trimester of pregnancy, low amniotic fluid levels serve as an important indicator of renal agenesis in the fetus [19]. Insufficient amniotic fluid, also known as oligohydramnios, in addition to the absence of kidneys, poses a significant risk for fetal death. In contrast, the impact of bilateral renal agenesis is markedly different in mice, resulting in death within 2 days after birth [20]. While the mouse phenotype we observed in our *Frem2-KO* mice is clearly more severe when compared to other previously reported *Frem2* mouse lines, given the uncertain rate of miscarriages related to FREM2 dysfunction, it is difficult to directly compare the severity of the phenotype we observed to that of patients carrying pathogenic FREM2 variants.

In summary, to address the gap in Fraser syndrome research, we have developed and characterized a constitutive *Frem2-KO* mouse model that lacks the FREM2 protein. While previous studies have utilized various *Frem2* mouse models, some of which have retained limited FREM2 function, here we report a mouse model predicted to lack FREM2 entirely. Our findings reveal that *Frem2-KO* pups die shortly after birth, limiting the opportunity for extensive study. Nevertheless, we have identified the most prominent phenotypes and the primary cause of neonatal mortality. We acknowledge the possibility of non-morphological phenotypes, particularly in the respiratory or cardiovascular systems, that have yet to be uncovered. Our study provides a unique, though severe, model of Fraser syndrome that can be utilized in future embryonic studies to advance the understanding of Fraser syndrome’s pathophysiology. Critically, our work can help tease apart the crucial role the FREM2 has in the development of important organs and also in uncovering the underlying mechanisms that drive Fraser syndrome.

## MATERIALS AND METHODS

### Animals

Mouse ES cell clones (*Frem2*^*tm1a(EUCOMM)Hmgu*^; clones HEPD0988-1-E06 and HEPD0988-1-H08) were purchased from the European Conditional Mouse Mutagenesis Program (https://www.mousephenotype.org/about-impc/about-ikmc/eucomm/, Clone# HEPD0988_1_E06, HEPD0988_1_H08). The clones were then used to generate the *Frem2*^*tm1a(EUCOMM)Hmgu*^ mouse line carrying the knockout-first allele with the conditional potential (F2KCR allele) (**Fig. 1A**) at the Harvard Transgenic Animal Core. For clone HEPD0988-1-E06, 36 blastocysts were injected, 8 pups were born (5 died), and no chimera pups obtained from this injection. For clone HEPD0988-1-H08, 48 blastocysts were injected, 15 pups were born (5 died), and two of the 10 surviving pups were chimeric mice (1 male was 15% and 1 female was 40% chimeric). The two chimera mice were bred to get germline transmission. The founder mice were then imported to the Mass Eye and Ear animal facility to set up an animal colony. To generate the conditional *Frem2*^*fl/fl*^ allele the FRT-flanked Neo cassette was excised by crossing with a FLP deleter strain (The Jackson Laboratory strain #003946).

The *knockout*-first mouse line (*Frem2*-*KO*-first allele with conditional potential, F2KCP) derived from these ES cells contains a cassette in the middle of exon 1 of the wildtype FREM2 gene. Upon further investigation, we determined that the sequence “GTCCCAGGTCCCGAAAACCAAAGAAGAAGAACGCA,” referred to as “exon 2” in the vendor’s documentation, does not align with any region of the published wild-type *Frem2* gene (Ensembl ENSMUSG00000037016.12). The F2KCR line was bred with the FLP mouse (JAX strain # 003946) to delete the “FRT-Exon2-LacZ-LoxP-NeoR” part and obtain a floxed FREM2 line containing “Exon1-FRT-LoxP-Exon3 (or Exon1b)-LoxP” structure.

All procedures and protocols were approved by the Institutional Animal Care and Use Committee of Mass Eye and Ear and were carried out in compliance with all relevant ethical regulations for animal use. All methods are reported in accordance with ARRIVE guidelines (https://arriveguidelines.org). All mice were kept on a 12:12 hour light-dark cycle with unlimited access to food and water. The fetuses were collected by timed breedings or by keeping track of the pregnancy by weighing the dam. For the collection of newborn pups, the breeding cages with pregnant females were closely monitored for litter birth by an infrared webcam setup below the cage with a remote access functionality.

### Phenotypic Analysis and Fixation of Fetuses

Mouse fetuses were collected by euthanizing pregnant adult females at the needed gestation stage. Fetuses were collected and fixed in Bouin’s fixative (Electron Microscopy Sciences). E12.6-16.5 were fixed for 4 hours. E17.5-18.5 were fixed for 72 hours. The fetuses were then rinsed in three changes of 70% alcohol and stored at 70% alcohol until processing.

### Collection and Fixation of Newborn Pups

Newborn mouse pups were collected at P0 once the litter was detected using an infrared webcam setup (ELP 1080p USB Camera). Newborn pups were anesthetized on ice, weighed, imaged, and the tip of the tail was collected for genotyping. Pups were then euthanized by decapitation and fixed in Bouin’s solution (Electron Microscopy Sciences, Cat# 15990-01) for 72 hours. After fixation, samples were rinsed in three changes of 70% alcohol and stored at 70% alcohol until processing. Pups used for gross anatomy evaluation were euthanized, and a median laparotomy was done from the pubic crest to the trachea.

### Histological Analysis

Fetal and newborn pup bodies were trimmed into three sections with cuts at the diaphragm and pelvis to ensure adequate paraffin embedding and tissue processing. Heads were trimmed for coronal embedding to the area of interest. Samples were processed using a Microm STP-120 processor on a routine program for manual paraffin embedding with a Microm EC-350 embedding center. After paraffin embedding, 5µm sections were cut from the head and body for Hematoxylin and eosin staining. Sections from the bodies were collected distally every ∼400 µm through the entire cavity. Slides were imaged with the Aperio AT2 automatic slide scanner (Leica Biosystems) equipped with Consolo v102.0.7.5 and Controller v102.0.8.704. The slides were digitized with a 20x/0.75NA Plan Apo objective lens and processed with the *Aperio ImageScope* software (version 12.4.6), available for download from https://www.leicabiosystems.com/us/digital-pathology/manage/aperio-imagescope/ free of charge. Anti-FREM2 immunolabeling was carried out using a previously reported labeling protocol[21]. The following antibodies against FREM2 were tested on the skin cryosectioned tissue samples of embryos: mouse monoclonal Cat. #SC-376555 (Santa Cruz Biotechnology); rabbit polyclonal Cat. #5831-1003 (ProScience) and rabbit polyclonal Cat. #5713, One World Lab (discontinued).

## Supporting information

Supplemental Movie 1 - Frem2-WT pup 1

Supplemental Movie 2 - Frem2-WT pup 2

Supplemental Movie 3 - Frem2-HET pup 1

Supplemental Movie 4 - Frem2-HET pup 2

Supplemental Movie 5 - Frem2-KO pup 1

Supplemental Movie 6 - Frem2-KO pup 2

## ACKNOWLEDGEMENTS

We thank Dr. David Corey for developing the *Frem2-KO* mouse line; Mr. Philip Seifert and the Schepens Eye Research Institute Morphology Core for sectioning eye samples for histological analysis; Ms. Sarah Visconti for assistance in animal breeding studies and histological imaging; Dr. Frank Yeh for editing and providing critical feedback on the manuscript. This work was supported by NIH R01DC020190 (NIDCD), R01DC021795 (NIDCD), and R01DC017166 (NIDCD) to A.A.I. The funder had no role in study design, data collection and analysis, decision to publish, or preparation of the manuscript.

## DATA AVAILABILITY

All data are included within the manuscript or are available from the corresponding author upon request. All imaging data will is available from a Zenodo public data repository https://doi.org/10.5281/zenodo.16649235.

## AUTHOR CONTRIBUTIONS

**RGS**: Validation, Formal Analysis, Investigation, Writing – Original Draft, Writing – Review & Editing, Visualization, Project administration.

**GZ**: Validation, Formal Analysis, Investigation, Writing – Review & Editing.

**OSS**: Validation, Formal Analysis, Investigation, Writing – Review & Editing.

**NL**: Investigation, Resources, Writing – Review & Editing.

**MZ**: Resources, Writing – Review & Editing, Visualization.

**AE**: Resources, Writing – Review & Editing.

**PYB**: Validation, Resources, Writing – Review & Editing.

**SW**: Conceptualization, Methodology, Validation, Formal Analysis, Investigation, Writing - Review & Editing, Visualization.

**LR**: Methodology, Validation, Investigation, Resources, Writing – Review & Editing, Supervision.

**AAI**: Conceptualization, Methodology, Validation, Formal analysis, Investigation, Resources, Visualization, Writing – Original Draft, Supervision, Project administration, Funding acquisition.

## SUPPLEMENTAL MATERIALS

**Supplemental Movie 1. Serial sectioning of newborn *Frem2***^***+/+***^ **pup 1**. Movie of images from hematoxylin and eosin-stained histology sections from a *Frem2*^*+/+*^ newborn pup. Transverse sections through the body begin inferior to the front paws and end superior to the hind paws displaying major organs including the lungs, heart, liver, intestines, the urinary bladder, and kidneys. Scale bar: 2 mm

**Supplemental Movie 2. Serial sectioning of newborn *Frem2***^***+/+***^ **pup 2**. Movie of images from hematoxylin and eosin-stained histology sections from a *Frem2*^*+/+*^ newborn pup. Transverse sections through the body begin inferior to the front paws and end superior to the hind paws displaying major organs including the lungs, heart, liver, intestines, the urinary bladder, and kidneys. Scale bar: 2 mm.

**Supplemental Movie 3. Serial sectioning of newborn *Frem2***^***+/-***^ **pup 1**. Movie of images from hematoxylin and eosin-stained histology sections from a *Frem2*^*+/-*^ newborn pup. Transverse sections through the body begin inferior to the front paws and end superior to the hind paws displaying major organs including the lungs, heart, liver, intestines, the urinary bladder, and kidneys. Scale bar: 2 mm.

**Supplemental Movie 4. Serial sectioning of newborn *Frem2***^***+/-***^ **pup 2**. Movie of images from hematoxylin and eosin-stained histology sections from a *Frem2*^*+/-*^ newborn pup. Transverse sections through the body begin inferior to the front paws and end superior to the hind paws displaying major organs including the lungs, heart, liver, intestines, the urinary bladder and kidneys. Scale bar: 2 mm.

**Supplemental Movie 5. Serial sectioning of newborn *Frem2***^***-/-***^ **pup 1**. Movie of images from hematoxylin and eosin-stained histology sections from a *Frem2*^*-/-*^ newborn pup. Transverse sections through the body begin inferior to the front paws and end superior to the hind paws displaying major organs. No urinary bladder can be easily identified and no kidneys are present. Scale bar: 2 mm.

**Supplemental Movie 6. Serial sectioning of newborn *Frem2***^***-/-***^ **pup 2**. Movie of images from hematoxylin and eosin-stained histology sections from a *Frem2*^*-/-*^ newborn pup. Transverse sections through the body begin inferior to the front paws and end superior to the hind paws, displaying all major organs. No urinary bladder can be easily identified and only the left kidney is present. Scale bar: 2 mm.

**Supplemental Figure 1.**
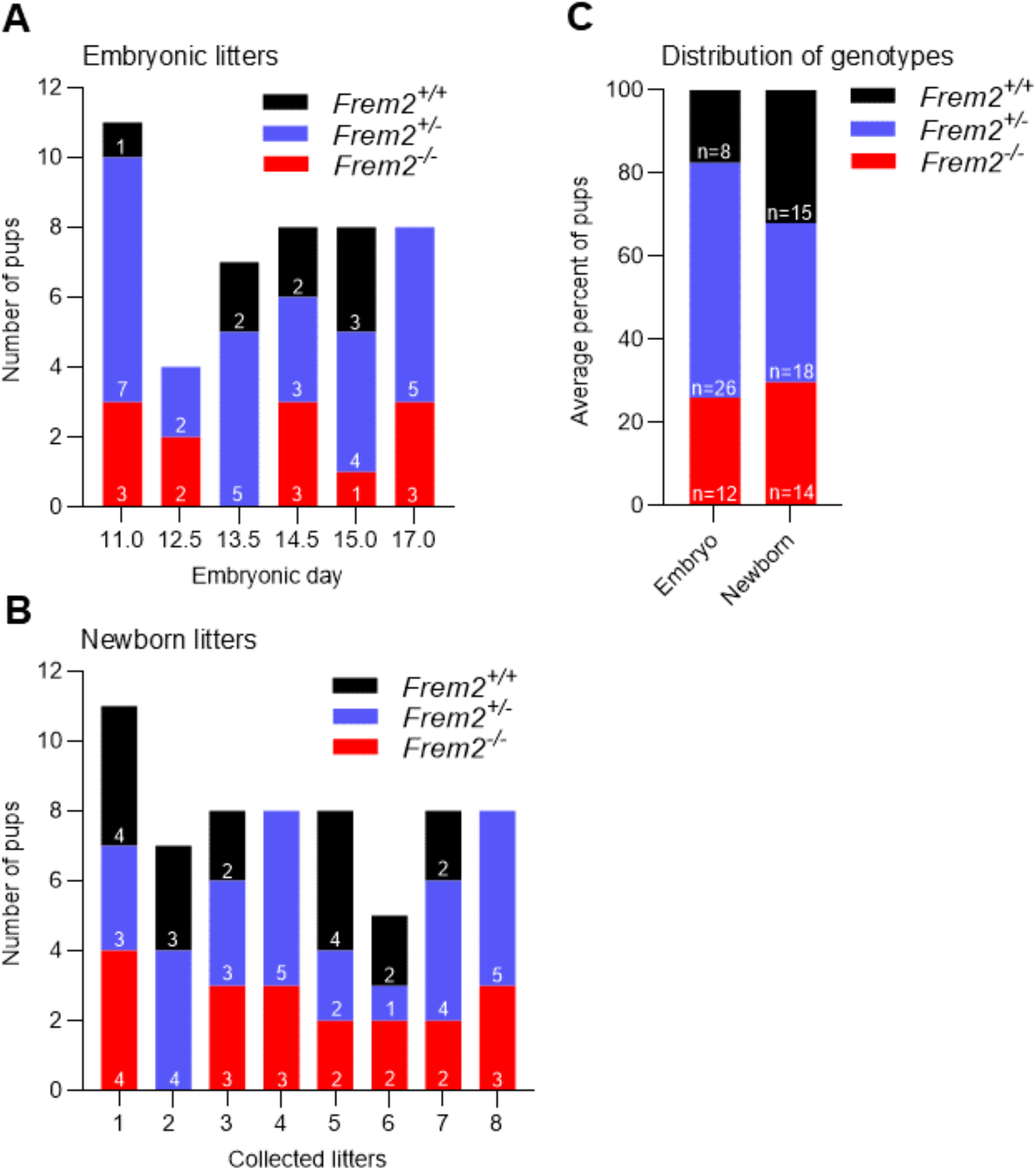
Genotype frequency in Frem2 litters. All litters were obtained from *Frem2*^*+/-*^ × *Frem2*^*+/-*^ crosses. (A) The number of *Frem2*^*+/+*^, *Frem2*^*+/-*^, and *Frem2*^*-/-*^ embryos collected from pregnant *Frem2*^*+/-*^ females at various gestational stages. (B) The number of *Frem2*^*+/+*^, *Frem2*^*+/-*^, and *Frem2*^*-/-*^ newborn pups in 8 litters from 7 different parents. (C) The distribution of genotypes across all embryonic (left) and newborn litters.

**Supplemental Table 1.**
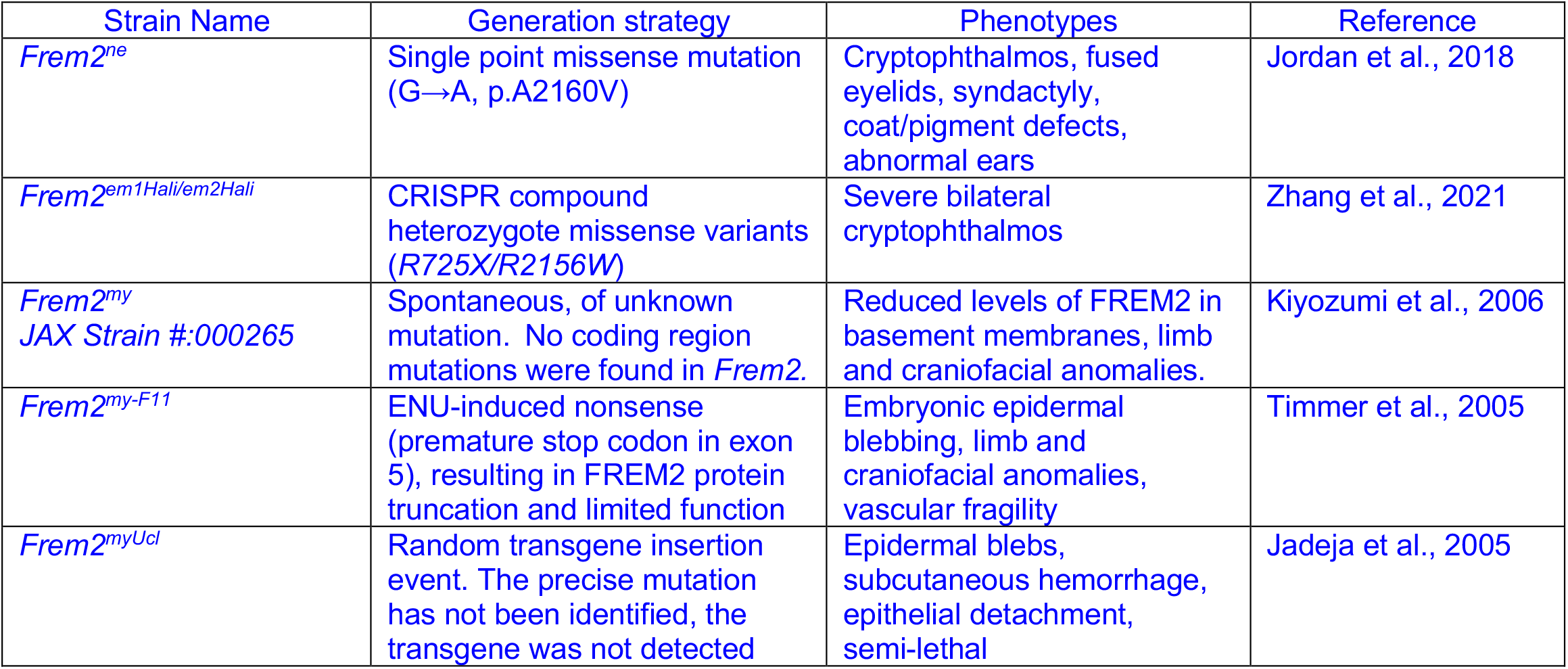
List of previously reported *Frem2* mouse strains.

**Supplemental Table 2.** Frequency of phenotypic presence in *Frem2-KO* embryos and newborn mice.

| Age | Number of *Frem2*-KO | Renal agenesis | Ocular hemorrhage | Soft-tissue syndactyly | Paw blister |
| --- | --- | --- | --- | --- | --- |
| E12.5 | 2 | not assessed | 0 | 2 | 2 |
| E13.5 | 3 | not assessed | 3 | 3 | 3 |
| E14.5 | 3 | not assessed | 3 | 3 | 3 |
| E15.0 | 1 | not assessed | 1 | 1 | 1 |
| E17.0 | 3 | not assessed | 3 | 3 | 3 |
| P0 | 3 | 3 | 3 | 3 | 2 |
| P0 | 4 | 4 | 4 | 4 | 3 |
| P0 | 2 | 2 | 2 | 2 | 2 |
| P0 | 3 | 3 | 3 | 3 | 3 |
| P0 | 2 | 2 | 1 | 1 | 1 |
| P0 | 2 | 2 | 2 | 2 | 2 |
| P0 | 2 | 2 | 2 | 2 | 2 |

For each embryonic and newborn litter collected, *Frem2*-KO animals were evaluated for prevalent phenotypes present such as renal agenesis, ocular hemorrhage, soft-tissue syndactyly, and paw blisters. The presence of kidneys was not evaluated in embryos.

